# CNSistent integration and feature extraction from somatic copy number profiles

**DOI:** 10.1101/2024.12.23.630118

**Authors:** Adam Streck, Roland F. Schwarz

## Abstract

The vast majority of cancers exhibit Somatic Copy Number Alterations (SCNAs)—gains and losses of variable regions of DNA. SCNAs can shape the phenotype of cancer cells, *e*.*g*. by increasing their proliferation rates, removing tumor suppressor genes, or immortalizing cells. While many SCNAs are unique to a patient, certain recurring patterns emerge as a result of shared selectional constraints or common mutational processes. To discover such patterns in a robust way, the size of the dataset is essential, which necessitates combining SCNA profiles from different cohorts, a non-trivial task.

To achieve this, we developed CNSistent, a Python package for imputation, filtering, consistent segmentation, feature extraction, and visualization of cancer copy number profiles from heterogeneous datasets. We demonstrate the utility of CNSistent by applying it to the publicly available TCGA, PCAWG, and TRACERx cohorts. We compare different segmentation and aggregation strategies on cancer type and subtype classification tasks using deep convolutional neural networks. We demonstrate an increase in accuracy over training on individual cohorts and efficient transfer learning between cohorts. Using integrated gradients we investigate lung cancer classification results, highlighting *SOX2* amplifications as the dominant copy number alteration in lung squamous cell carcinoma.

## Introduction

Somatic copy number alterations (SCNAs)—gains and losses of long regions of DNA—are found across almost all cancer types and are one of the key defining features separating cancer cells from normal cells^1^. It has been demonstrated that quantifying SCNAs has predictive value in the clinic for both progression free and overall survival ^2,3^ and that they can serve as sensitive biomarkers for cancer classification and subtyping^4^. We and others have shown that many cancers demonstrate ongoing chromosomal instability (CIN) and continuously accumulate SCNAs throughout their evolution^5^, and that SCNAs are excellent markers for inferring cancer evolution^6,7^.

SCNA profiles are commonly derived from a variety of experimental techniques, including SNP arrays, whole-exome and whole-genome sequencing^8^, and recently also increasingly from single-cell sequencing^9^. One major advantage of SCNAs over other genomic data types including somatic single nucleotide variants (SNVs) is ease of handling. Due to their aggregate nature, SCNA profiles of individual patients can be published without concern for privacy and the resulting access restrictions, leading to a growing set of publicly available and easily accessible samples from large cohorts such as TCGA, ICGC^10^, and the TRACERx^11^ lung and renal cancer cohorts.

Now with more than 15000 genomic profiles available in these cohorts alone, sophisticated statistical and machine learning algorithms for understanding CIN and the origins of SCNAs are being developed. For example, copy number signatures have linked SCNAs to their underlying molecular mechanisms, further strengthening their prognostic value^12,13^. However, most large-scale studies, including the ones just mentioned, still rely on profiles from single cohorts such as TCGA. Between cohorts, SCNA profiles are more difficult to integrate. Different experimental techniques and different copy number calling algorithms can lead to downstream biases, and varying granularities, bin sizes or segment lengths further complicate a joint analysis of SCNA profiles from different cohorts.

In order to fully leverage these profiles, data scientists require a unified method for processing and integration of these datasets. Deep learning in particular requires big, clean datasets, even more so as the size of models increases with the potential goal of building foundational models^14^. Since reprocessing several thousand whole genomes is often prohibitive due to computational costs and access restrictions, methods to integrate and unify publicly available SCNA profiles are highly useful.

We here present CNSistent, a Python package for preprocessing, consistent segmentation, integration, statistical analysis and visualization of SCNA profiles coming from heterogeneous data sources. We demonstrate the utility of CNSistent by integrating available copy-number profiles from the TCGA, PCAWG and TRACERx cohorts. Once processed, we evaluate various segmentation strategies, comparing the performance of deep learning-based multiclass cancer classification tasks and on the classification of non-small cell lung carcinomas (NSCLC). Our method outperforms the previously published results in both cases, in the latter case quite significantly and we also demonstrate that a model trained on one dataset generalizes well to other datasets. Lastly, we use explainable AI methods to discover regions of interest in the NSCLC datasets.

## Methods

CNSistent processes SCNA profiles using a multi-step approach. Input data takes the form of copy number segment tables with either allele-specific or total copy numbers (Fig. 1A) for which CNSistent calculates the proportion of missing segments. Optionally, masking can be used in this or the following steps (Fig. 1B) to blacklist genomic regions of low interest. CNSistent then utilises imputation strategies to fill in missing data (Fig. 1C) and calculates statistical features for each SCNA profile (Fig. 1D). CNSistent then offers various strategies for creating a consistent segmentation across samples (Fig. 1E), which are subsequently aggregated to create a final set of complete SCNA profiles with shared segment boundaries for all samples (Fig. 1F). Each of these steps is detailed in its own section below. The pipeline is fully modular and the steps can be skipped or executed out of order.

**Figure 1:**
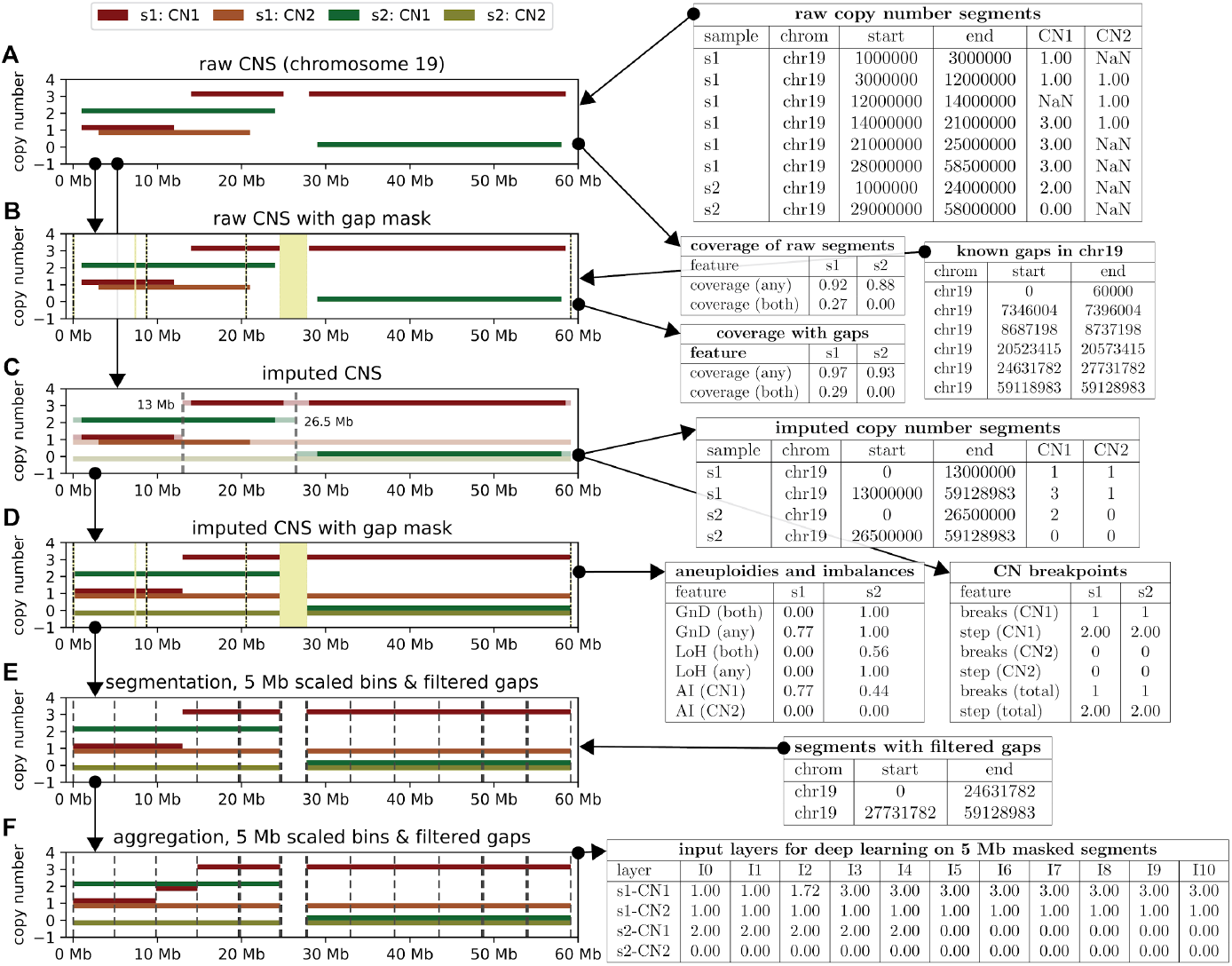
Illustrative example of processing of two SCNA profiles (s1, s2) for two alleles (CN1, CN2), on human chromosome 19. A) The input data consist of non-contiguous major and minor copy number segments for each sample. B) Calculation of the **proportion of the genome that is missing** for each sample. For comparison, the coverage is calculated both with and without considering the blacklisted regions. Note that as there are no minor CNs for s2, the homozygous coverage is 0. C) During **imputation** two new breakpoints are introduced at 13 Mb and 26.5 Mb, while the breakpoints on the boundaries of missing segments are no longer present. From the imputed data CNSistent calculates the CN **breakpoint-related statistical features**. D) Ploidy-related statistical features that are derived from the imputed data and removal of the blacklisted regions. E) Small regions are removed from masking, retaining only the gap between 20 and 30 Mb, which splits the chromosome into two arms, these are then further split into ~5 Mb bins. The same-size strategy is used, meaning that the bins in the left segment are slightly smaller (4.9 Mb), while the ones on the right are slightly bigger (5.27 Mb). F) Aggregation of CNs across samples and alleles. This is then converted into vectors of input values for downstream deep learning, each value representing one input node. Note that as there was a breakpoint at 13 Mb, the resulting value is a weighted mean of the previous values, i.e. 1.72.

For its calculations, CNSistent can work with any reference genome; hg19 and hg38 reference assemblies are provided as a default. If the sex of the donors is not provided, CNSistent will determine the sex for each sample based on the presence of the Y chromosome.

### Segment imputation

SCNA profiles from different cohorts often vary in the extent to which they span the genome. This can be due to a variety of reasons including different underlying technologies (WES vs WGS sequencing), different segmentation strategies, or different blacklisting of regions surrounding the centromeres and telomeres. To retain as much information as possible, CNSistent offers an imputation step capable of filling the gaps in SCNA profiles using an *extension* method (Fig. 1C).

The extension imputation method executes the following five steps: (i) Segments are pruned such that they are fully contained within the coordinates and named chromosomes of the reference genome. (ii) CNSistent extends the first and last segment of each chromosome to the chromosome boundaries. (iii) Each gap between two segments is split into two halves (rounded down), and each half is then assigned the CN of its neighboring segment. (iv) If any chromosomes are fully missing from the sample, they are set to 0. (v) The neighboring segments that have the same CN are merged.

Alternatively, two additional imputation options are available: *diploid* and *null*. The diploid method changes the steps (ii-iv) in such a way that all newly created segments are set to diploid, e.g. if a sample is male and major/minor CN columns are used, CNSistent will create a segment on the whole chromosome Y with major and minor CN of 1 and 0 respectively. The null option will analogously fill all the newly created segments with 0.

### Feature extraction

CNSistent can calculate a set of statistical features. As CNSistent is sex chromosome aware, the length of the linear genome depends on the sex of the sample. Each of the following features is therefore calculated three times: for autosomes, for sex chromosomes, and for the whole genome:

#### Coverage

Calculates the proportion of the whole genome where any CN value is assigned (as opposed to missing values). In case of allele-specific CNs, both mono-allelic coverage (either allele has a CN value assigned) and bi-allelic coverage (both alleles must have a CN value assigned) are calculated.

#### Genome not Diploid (GnD)

Defines the proportion of the genome where an allele does not have the CN a diploid cell of the same sex would have. In case of having only total CN this is a lower bound approximation.

#### Loss of Heterozygosity (LoH)

Calculates the proportion of segments with CN=0 on either allele (hemizygous) or on both alleles (nullizygous). The segment is only considered LoH if and only if its CN value is 0 and its normal value is not zero (e.g. chromosome Y for female).

#### Allelic imbalance (AI)

The proportion where one allele has a strictly higher CN than the other.

#### Breakpoints

The number of breakpoints per chromosome for each allele. If two column format is used, a total number of breakpoints is also calculated to account for cases where both alleles have a breakpoint in the same location (meaning that the total number of breakpoints is less than the sum of the alleles).

#### Breakpoint Step

The mean difference between the CNs of consecutive segments. Note that it is preferable to impute the segments first to avoid inducing spurious gaps.

### Consistent segmentation

One major goal of CNSistent is to obtain segments that are consistent between sample sets and from which then features can be derived in a unified manner. This requires the same set of breakpoints to be present in every sample. Segmentation consists of the following 4 steps: (i) create regions of interest, (ii) remove blacklisted regions, (iii) merge existing breakpoints (iv) subdivide the segments based on size. Each of the four steps is optional.

The segments for step (i) can be provided as a BED file, or one of five predefined options can be used: whole chromosomes (default option), chromosome arms, cytobands, COSMIC consensus cancer gene set^15^, or the Ensembl coding genes set^16^. From these segments, blacklisted regions can be optionally removed (Fig. 1D). As a default option, the regions of low mappability as defined by the UCSC^17^ genome browser are provided. During the blacklisting, if the regions are small or close to each other, fragmentation can occur. This can be avoided by segment filtering—the user specifies a filter of size f, where any blacklisted region smaller than f is removed, and likewise if after the blacklisted regions are removed from segmentation, any newly created segments smaller than f are also removed.

The breakpoints are then merged using a greedy algorithm on a predefined region (usually a whole chromosome). Starting from the leftmost breakpoint, all breakpoints within the merge distance ***m*** are accumulated and a new breakpoint is created as their average. This is then repeated from the leftmost not yet merged breakpoint, until the end of the region is reached. A detailed example is shown in Supp. Fig. 1.

Lastly, the resulting segments can be subdivided into smaller bins based on user defined split size ***s*** (Fig. 1E). Three subdivision strategies are provided: (a) From the start of the segment, breakpoints are inserted every ***s*** bases. Here, the last bin is likely to be of a different size. If it is smaller than ***s/2***, it is merged with the previous segment. (b) Is similar to (a) where instead of creating the padding only at the end, the padding is split in half and added to both ends. Likewise, if the first and last bins are smaller than ***s/2***, they are merged with their neighboring segments. (c) The bins are scaled so that they are all the same length, slightly different from ***s***. Consider a segment that has ***c*** bins, including the padding–If the padding is smaller or equal to ***s/2***, split the segment into ***c-1*** equally sized bins, otherwise into ***c*** bins.

### Aggregation of copy numbers

After joint segmentation, the copy numbers from the original segments are aggregated to create CNs for the new segments. First the old segments are split at the breakpoints given by the new segmentation. Second, the resulting refined segments are aggregated between the breakpoints given by the segmentation, using one of four possible aggregation strategies: The *Min* and *Max* strategies will assign the minimum or maximum CN to the whole segment –the min strategy is particularly relevant when considering genes, since incomplete segments are unlikely to yield functional gene copies. The *Mean* strategy will take a mean of CNs across bins weighted by their lengths, preserving the overall CN per sample. Lastly, merging can be skipped altogether, which can be used if we want to select only a subsection of each profile, e.g. only q-arms.

### Sample filtering

The features obtained in the feature extraction step can be used to filter undesirable samples. For base quality metrics, like coverage, a simple z-score outlier detection method is provided, meaning that for a feature *f* over a set of samples 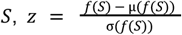 is calculated and samples greater than 3 standard deviations from the mean (|*z*| ≥ 3) are removed. The value 3 is a typical threshold for the method, but it can be adjusted by the user.

In certain cases, a qualitative separation of data is preferable, e.g. to remove samples with negligible SCNA activity. CNSistent offers an automated solution for finding such thresholds using a knee-detection algorithm. A knee-point is a point of the plot where the maximum angle between the line to the first point and the last point of the plot. To find the knee-points for a feature *f* in a set of samples *S*, a tuple of monotonically increasing feature values *T* = (*min*(*f*(*S*), …, *max*(*f*(*S*)), ∀*i* ∈ 1, |*T*| − 1: *t*_*i*_ ≤ *t*_*i*+1_ and a cumulative distribution of values smaller than each threshold *Y* = (|*f*(*S*) ≤ *t*|)_{*t* ∈*T*}_ is created. Second, *T* is normalized such that 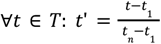 and analogously for *Y*′. The knee-point is then the *t*_*i*_, 1 ≤ *i* ≤ *n* with the maximum angle between the vector from origin to the normalized threshold, (*t*_*i*_ *′, y*_*i*_ *′*), and the vector from the threshold to the endpoint, (1 − *t*_*i*_ ′, 1 − *y*_*i*_ ′). If the angle is negative (clockwise rotation), we call it a *knee*, otherwise (counter-clockwise rotation), we call it an *elbow*.

### Deep Learning classification and attribution

For the classification tasks used to benchmark CNSistent, we used a convolutional neural network (CNN) with an architecture roughly similar to the one described in Attique et al^18^. The model has two padded convolutional (kernel=5) and maxpool (kernel=4) layers, followed by a fully connected layer with dropout (p=0.8) and then an output layer with size equal to the number of classes (see Supp. Fig. 3 for a visual schema). Each of the two input alleles is assigned one input channel. Different segmentation sizes are accommodated using a width-adjustable CNN^19^ with variable input layer width. The size of the output is likewise scaled w.r.t. the number of classes under classification.

The resulting accuracy is validated using 5-fold cross-validation, i.e. the dataset is separated into 5 equally sized subsets with a 4:1 train-test ratio. The resulting accuracies are the mean of the 5 training runs. As the number of patients per cancer type varies, the classes are imbalanced. To avoid a possible bias due to an overrepresentation of one class, stratified split is used, meaning that the ratio of the individual cancer classes is preserved across the 5 subsets. Additionally, some samples are obtained through multi-region sampling. While the samples from different regions show different profiles, there is a risk of being able to guess the class based on the similarity to the original profile. This is prevented by sample grouping where each group (in this case patient) can only be part of one subset. The splitting is done using the StratifiedGroupKFold object from scikit-learn v1.4.1.

For training we used the PyTorch 2 library^20^, torch package v2.2.1, accelerated using CUDA v12.1. Optimization was conducted using the Adam optimizer with a learning rate of 0.001 and weight decay of 0.01. The error is evaluated using cross-entropy loss. The training was limited to 1000 epochs. The training process was accelerated by an early stopping strategy where the minimum loss is recorded and if past 10 epochs lead to a training loss higher than the existing global minimum, the training stops.

To observe which segments are instrumental in determining the type of cancer during classification, we used the integrated gradients (IGs)^21^. These were computed using the package Captum v0.7.0^22^ with 20 approximation steps.

### Data and Code availability

CNSistent package, source data, and plotting notebooks can be downloaded at: https://bitbucket.org/schwarzlab/cnsistent.

The data produced by CNSistent are available at https://zenodo.org/records/14547456 ^23^.

The deep learning code and results are available at https://zenodo.org/records/14546762 ^24^.

## Results

We illustrate the use of CNSistent on a cancer type classification task using 15,072 publicly available SCNA profiles from The Cancer Genome Atlas (TCGA^13^, n=10674), the Pan-Cancer Analysis of Whole Genomes (PCAWG^10^, n=2778) and the TRACERx cohort of non-small cell lung cancer^11^ (n=1620).

Where the TCGA and PCAWG datasets overlap (829 samples), we gave preference to the PCAWG callset. The PCAWG dataset blacklists 195 low-quality samples, which were removed before further processing. The TRACERx dataset consists of two parts, primary tumor samples (n=1428) and primary with metastatic samples (n=694). We used the primary sample set in the 502 samples on which they overlap. This yielded a total set of 14174 SCNA profiles which were subjected to CNSistent for pre-processing and integration. A summary of sample counts is provided in Supp. Table 1.

### Feature distributions vary across datasets

We started by imputing any missing data and calculated the sample features (see Supp. Fig. 4 for complete results). Since SCNA profiles for sex chromosomes were not available in the TRACERx cohort, all sex chromosomes were removed from further analysis. Before masking, the SCNA profiles covered on average 98.47%, 96.39%, and 91.18% for PCAWG, TCGA and TRACERx respectively (Fig. 2A). When blacklisting using the UCSC gap regions, the coverage rose to 99.62%, 99.89%, and 97.38%. The gap regions of hg19 on autosomes sum to 19.65 Mb, which is 6.82% of the total genome. For TCGA and PCAWG virtually all the missing segments fell into these blacklisted regions. In TRACERx, there are regions missing also outside these gap regions, however mostly on their boundaries (Fig. 2C). This was likely due to the sequencing method: PCAWG data has been sourced using WGS and TCGA combines multiple data sources.

**Figure 2:**
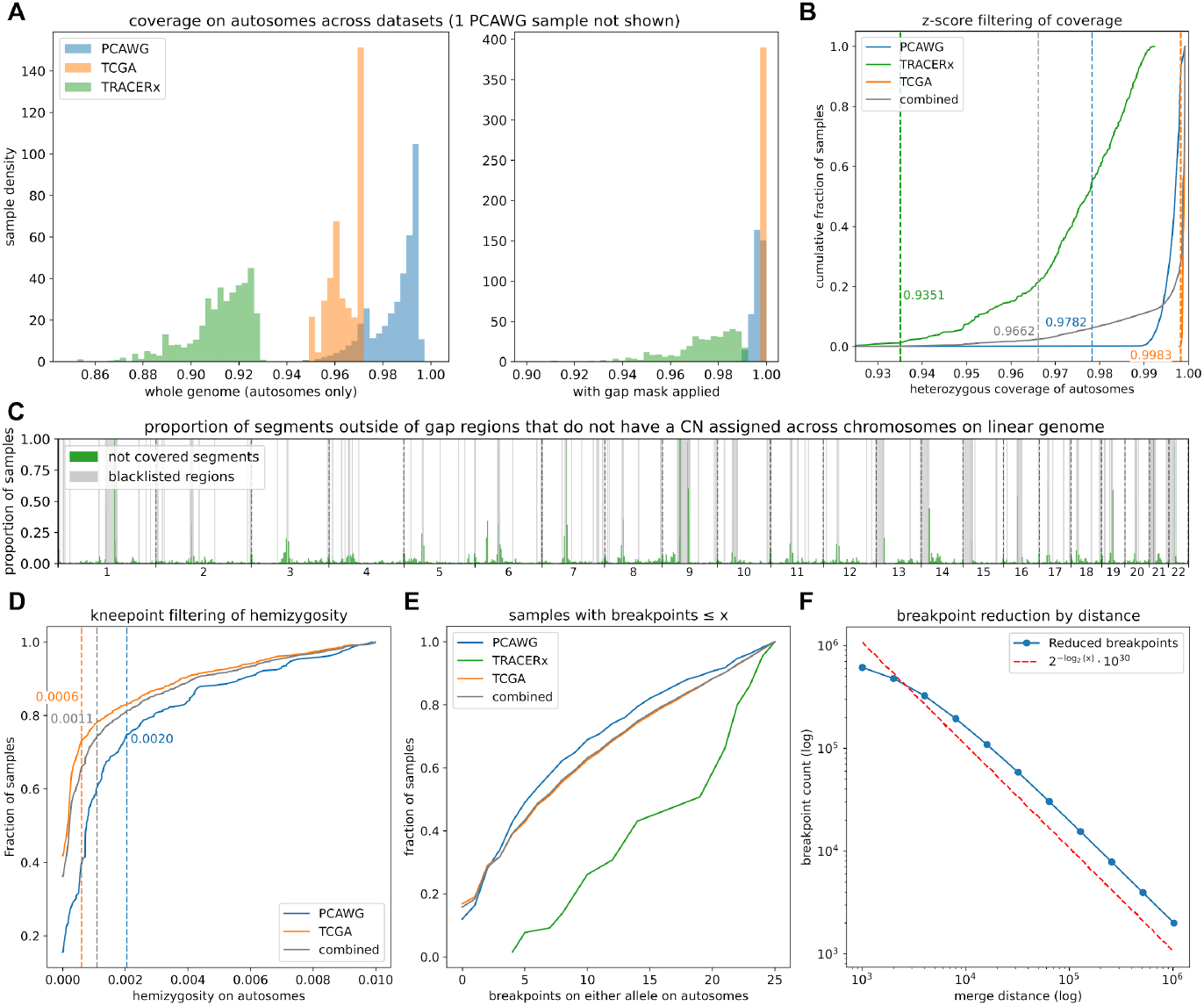
Processing of the PCAWG, TRACERx, and TCGA datasets. **A)** Histograms of heterozygous coverage before and after gap region filtering. Note that the PCAWG and TCGA datasets have almost full coverage after filtering. In contrast, while TRACERx shows a major shift, there are still substantial portions missing. **B)** Cumulative distribution of samples by heterozygous coverage, with the threshold for filtering given by the z-score. The position of the threshold is much higher for the combined dataset compared to the individual ones. **C)** Distribution of the missing values in the TRACERx dataset along the linear chromosomes. Data are mostly missing in regions close to the centromeres and telomeres, in particular for chromosomes 1 and 9. The TRACERx set is missing the sex chromosomes, therefore the X and Y are either declared as missing (green) or gap (gray) in all the samples. **D)** Cumulative distribution of GND for a subset of samples below 1%. TRACERx is not shown as none of the samples has hemizygosity below this value. Note the clear slope change around 0.1%, also detected by our kneepoint algorithm. **E)** Cumulative distribution of breakpoint counts for a subset of samples with less than or equal to 25 breakpoints. The curve is almost linear for all datasets, demonstrating that there’s no clear cutoff value in this region. **F)** The result of breakpoint reduction using 11 log-distributed merge distances between 1 Kb and 1 Mb. Note that the relationship is proportional—doubling the distance leads to halving the number of resulting segments, as shown by the hyperbolic curve.

Next, samples with low coverage were removed using the z-score based outlier detection (Methods). Thresholds were calculated for each of the datasets separately as well as using the combined dataset of all samples (Fig. 2B). For the individual samples there was only a small set of outliers: 3, 16, and 19 for the thresholds of 97.82% for PCAWG, 99.83% for TCGA, and 93.51% for TRACERx respectively. However, when the combined dataset was used, 352 samples were below the detected threshold of 96.62%, stemming from the fact that the coverage distribution of TRACERx significantly differs from the other two. In this case, filtering each set separately leads to significantly lower removal rate. Additionally, one sample in the PCAWG dataset, SP107557, had coverage of only 57.67% and presumably should have been blacklisted in the original dataset.

As we were interested in identifying cancer types from SCNA profiles, we also do not want to consider samples that have very few alterations, as these do not hold sufficient information for the classification task. Other authors have used the number of breakpoints^13^ as evidence for SCNAs, however we have not observed a clear knee-point in the data (Fig. 2E), and used the GnD proportion instead. We first set a threshold at 1%, meaning that any sample with over 1% of GnD was retained. In TRACERx, all samples were above this threshold, however in PCAWG and TCGA, ~10% of samples were below. To further increase the retention of samples, we looked for a qualitative change in the sample below 1%. In Fig. 2D one can observe a significant change in the slope around 0.1%, meaning that there is an accumulation of samples below this value, which we wanted to remove. To obtain an exact threshold, we used our knee point detection algorithm, obtaining the threshold of 0.06% for TCGA (745 samples), 0.2% for PCAWG (211 samples), and 0.11% for the combined one (965 samples). We used individual thresholds to filter the samples, and observed that the proportion of samples removed was quite similar: 8.16% of the total PCAWG set and 7.47% of the total TCGA set. The filtering process then leads to the final filtered sample set of 12901 samples (Supp. Fig. 5).

### One megabase segmentation provides the best variance/bias trade-off

We next set out to explore the effects of different segmentation strategies on the cancer classification task. CNSistent reported that over 50% of the samples have less than 50 breakpoints, and over 99% have less than 500 (Supp. Fig. 4D), and we therefore hypothesized that the optimal segmentation is likely within an order of magnitude of that. We thus processed the data using the following segmentation strategies: (i) fixed-size segments of 10 Mb, 5 Mb, 3 Mb, 2 Mb, 1 Mb, 500 Kb, and 250 Kb size; (ii) Whole chromosome, chromosome arm, and cytoband-level CN segments; (iii) Gene-level CN values based on the ENSEMBL and COSMIC gene sets; and (iv) breakpoint clustering using distance thresholds of 1 Mb, 500 Kb, and 250 Kb. To select the distance thresholds, we first evaluated the effect of varying the distance threshold on the resulting breakpoint count. Without any merging, the whole filtered dataset has 826910 unique breakpoints, i.e. one breakpoint per 3.7 Kb on average. Testing 11 merge distances from 1 Kb to 1 Mb we observed that the 1 Kb distance leads to 24.39% reduction, while 1 Mb leads to 99.65%, with only 2864 of the breakpoints left. As shown in Fig. 2F, after around 4 Kb, there is an almost perfect inverse relationship where doubling the merging distance leads to halving of the number of the resulting breakpoints. To match the other strategies in terms of segment count, we selected 1 Mb, 500 Kb, and 250 Kb distances, leading to 2797, 5569 and 10797 autosomal segments respectively. To compute all combinations of segmentation strategy and datasets efficiently, we made use of CNSistent’s internal parallelization strategy. Runtime decreased in a near-linearly with the number of compute cores available (Supp. Fig. 6). All segmentation configurations are listed in Supp. Table 2.

We subjected the resulting segmented datasets to the cancer type classification task. To our knowledge the best result to date has been reported on classification of top 6 cancer types in the dataset in Attique *et al*.^*18*^, with up to 92% test accuracy on the best model. Using our combined dataset, the selection of top 6 classes resulted in a set of 5172 samples with the following class labels: lung adenocarcinoma (LUAD, n=1314), breast invasive carcinoma (BRCA, n=1157), lung squamous cell carcinoma (LUSC, n=996), ovarian cancer (OV, n=618), prostate adenocarcinoma (PRAD, n=563), and kidney renal cell carcinoma (KIRC, n=513).

Training the 6-class classifier (Fig. 3A) test accuracy was 72.48% for the whole chromosomes, i.e. 2 channels with 22 input nodes each. After this point, the accuracy quickly increased with increasing granularity. However, at around 10^3^ segments test accuracy almost plateaus, while training accuracy keeps increasing, indicative of overfitting. This is visible in particular when comparing the COSMIC gene set (90.27% accuracy using 681 segments) to the Ensembl gene set (89.08% accuracy using 19430 segments).

**Figure 3:**
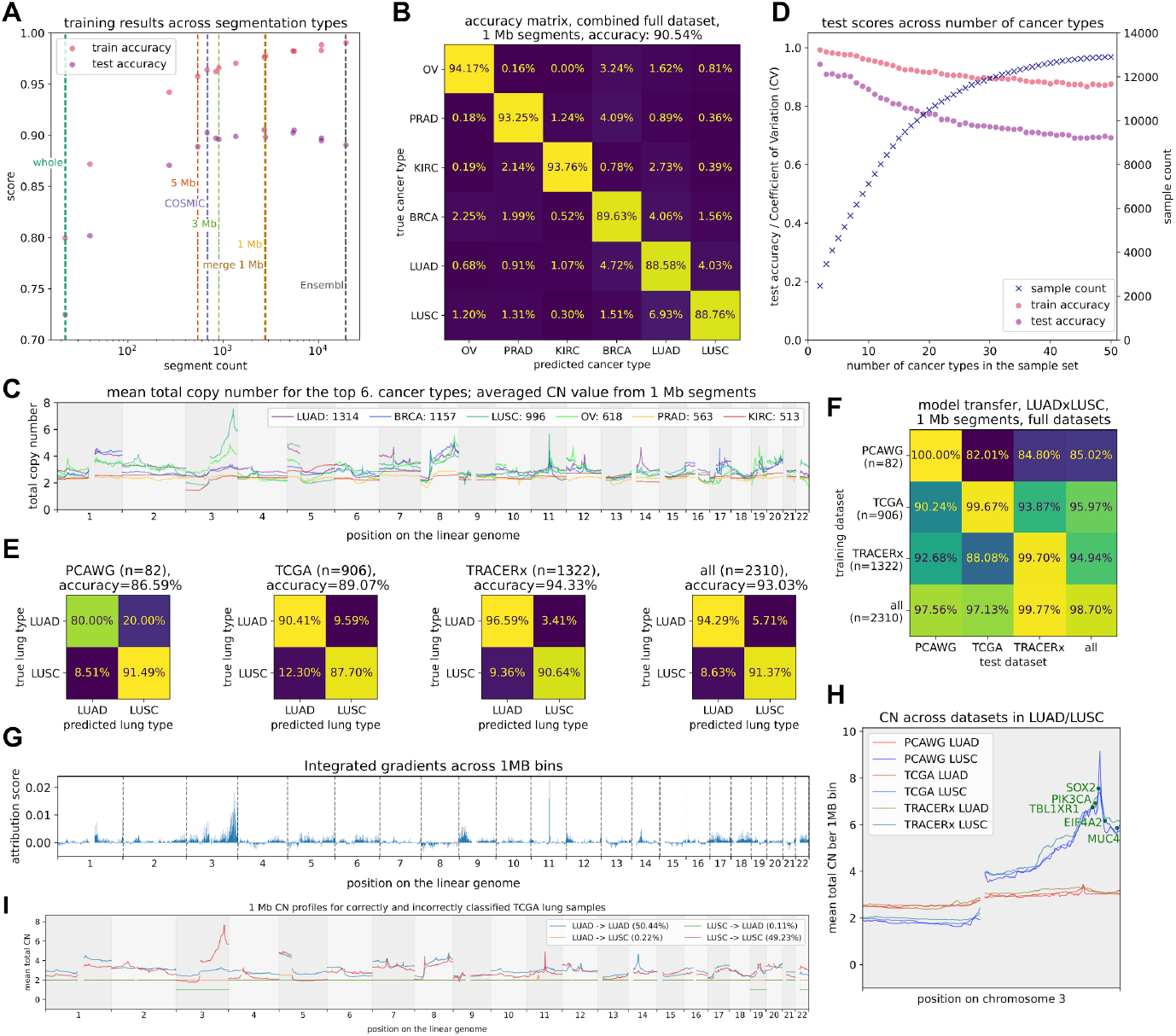
Evaluation of multi-class prediction task. **A)** Results of training across 15 different segment types, using two features (one for each allele) per segment. We see that around 10^3^ segments, the test accuracy starts to plateau and even decline, while training accuracy continuously increases, suggesting that more refined segmentation leads to overfitting. **B)** Confusion matrix for the best observed accuracy with 1 Mb segmentation. **C)** Copy number profiles of the sample set for 6-class classification using the 1 Mb segments. Sex chromosomes are not shown as they are not used in training. Each segment is represented at its centerpoint, thus the lines represent linear interpolations between segments. **D)** Training of classifiers for up to 40 classes. The samples are accumulating at decreasing intervals. Both the training and testing accuracies decrease in linear fashion, with the slope being steeper for the testing accuracies, presumably driven by the decreasing number of samples. **E)** Confusion matrices for the lung cancer classification for the PCAWG, TRACERx, TCGA, datasets and all three combined. The TRACERx sample set shows by far the best accuracy. **F)** The accuracy of testing on one dataset after training on another. The diagonal therefore also represents the training accuracy when training on all available samples. **G)** Integrated gradients over 1 Mb bins, demonstrating a strong focus on chromosome 3. **H)** CN plots for both LUAD and LUSC samples from each dataset. Notice a strong correlation between the LUAD sets (red tint) and LUSC sets (blue tint). The 5 genes with the highest attributions from IGs have CNs averaged from all samples. **I)** Plot of true and false positives in training on the TCGA dataset with 1 Mb bins, notice the similarity of chromosome 3 pattern for the correctly classified LUSC samples (red) and the misclassified ones (orange).

Seemingly, considering gene-coding regions only leads to a better performance than using non-coding ones—COSMIC set (681 segments, accuracy 90.26%) performs better than 5 Mb set (542 segments, 88.89% accuracy) and 3 Mb set (902 segments, 89.61% accuracy). Otherwise, we did not find any considerable difference between breakpoint merging and pre-defined segmentation. Similarly, using cytobands (834 segments, accuracy 89.72%) yielded almost the exact same result as the similarly sized 3 Mb segments. Full training results are given in Supp. Table 3.

### Improved accuracy over competing models and techniques

The best test cross-validated accuracy of 90.54% was achieved using 2709 1 Mb segments. In Attique *et al*.^*18*^ the authors report an optimal accuracy of 92% without cross validation. In our case the best test accuracy without cross-validation was 92.34% and the best test accuracy during training was 93.31%, rendering our model at least on par with the existing model. In the confusion matrix (Fig. 3B) one can observe that all the classes are predicted well, with the difference of the highest accuracy (PRAD at 94.17%) and the lowest (LUAD at 88.58%) being only 5.41%.

To better understand the structure of the classification task, we visualized the CN plots for the top 6 classes (Fig. 3C), which revealed that the patterns were quite distinct, with some well-known, prominent peaks, e.g. in chromosome 3 for LUSC^25^ and OV, or chromosome 14 for LUAD^26^. We have fitted a UMAP (Supp. Fig. 7) and observed there is no clear separation of clusters between cancer types. For comparison we also trained a Random Forest classifier (Supp. Fig. 8), which only yielded a 5-fold cross-validation accuracy of 82.52%, performing substantially worse (-8.02%) than the CNN.

Additionally we were interested in how our network performs for different numbers of classes. In Fig. 3D it can be seen that the accuracy is quite high for all the cases and decreases in almost a linear fashion. In the easiest 2-class task we saw 94.34% test accuracy, while the 50-class task reached 69.24%.

### Model transfer between datasets

We next tested each model on the other datasets (Fig. 3F). The PCAWG model does not transfer effectively (82.01% on TCGA and 84.80% on TRACERx), likely stemming from only having 82 lung cancer samples in total. Surprisingly, the TCGA-trained model has a higher accuracy when applied to the other two sets (90.24% and 93.87%) than when training on itself (89.07% with 5-fold cross-validation). The TRACERx model also transfers quite well, yielding a significantly higher accuracy on the PCAWG dataset (92.68%), compared to the PCAWG-trained model (86.59%). We therefore assumed that the TRACERx model does generalize well to unseen data and does not suffer from overfitting to match individual patients, even when multiple-region sampling is used to construct the dataset. Moreover, we observed that increasing the size of the training set leads to improvement in accuracy.

### Explainability and the effect of *SOX2* gene

To further investigate why the model trained on the TCGA dataset performs better on other datasets, we conducted explainability analysis using the Integrated Gradients method (IG)^21^. This has been shown to be an efficient method of identification of significant genes^27^. However, the authors therein were unable to investigate NSCLC subtypes due to missing labels which fortunately are present in our data.

First, we first calculated IGs on 1 Mb bins to obtain top-level understanding of the focus areas. Briefly, The IG method computes gradients between input and output nodes, attributing scores based on their influence. An increase in the output value for a specific class due to a high input value results in a positive attribution, while a decrease yields a negative attribution. The magnitude of the score reflects the input node’s importance to the output. Using IG we found chr 3 to play a major role in distinguishing between LUAD and LUSC (Fig. 3G). We therefore analyzed the patterns of CNs on this chromosome more closely (Fig. 3H). When considering the average CNs per dataset each dataset is surprisingly well correlated to the other two. In particular for LUSC, the Pearson correlation coefficient between PCAWG and TCGA is 0.9657. In the TRACERx, the correlation is not as strong, but still significant at 0.8851 against PCAWG and 0.8926 against TCGA, showing a systematic positive selection for q arm of chromosome 3 in LUSC.

To investigate the genes involved, we applied the IG method to the COSMIC gene set. The top 9 attributions were all to genes on the q arm of chromosome 3. We plotted the top 5 (Fig. 3B), each of which are known for NSCLC associations: *EIF4A2*^*28*^, *SOX2*^*29*^, *TBL1XR1*^*30*^, *MUC4*^*31*^, and *PIK3CA*^*32*^. When overlaid with the CN plot, it became evident that *SOX2* forms the peak of the profile, with the average CN=7.56. The 4 following highest CN values were *PI3CA* (6.91), *TBL1XR1* (6.75), *MAP3K13* (6.53), and *IGF2BP2* (6.44), which are the 2 closest left- and right-side neighbors of *SOX2* in the COSMIC dataset, suggesting that these are co-selected by the selection of *SOX2*. After *SOX2* there is a sharp drop in average CN, which plateaus around *EIF4A2* (average CN 6.16). We therefore propose that in LUSC there is a strong selection for *SOX2* combined with a selection against *EIF4A2*. Selection for *SOX2*^*29*,*33*^ is well-known, however poor prognosis has been associated both with upregulated^34^ and downregulated^28^ *EIF4A2*.

The only gene in top 10 attributions was *FADD* on chromosome 11, which exhibits significant local amplification in LUSC, also a known actor in lung cancer^35^, with the average CN of 5.64 (see Supp. Fig. 9 for details).

### Misclassification might be caused by insufficient labelling

We next used the IG results to investigate the mis-classification observed in TCGA. We used the results of training without cross-validation, i.e. all the samples post-filtering (908) were placed in the training set, resulting in the training accuracy of 98.46%. In the CN plots of the misclassified samples (Fig. 3I) there is one sample mis-classified as LUAD, which has no other pattern than loss of whole chromosomes 3, 19, and 22. This is likely to be difficult to classify using a CNN as there are no changes within chromosomes. Next, there were 13 samples mis-classified as LUSC. These samples exhibited a pattern of monotonically increasing gains on the q arm of chromosome 3, which we earlier associated with LUSC. Unlike in TRACERx, in the TCGA dataset there is no classification for samples having both LUSC and LUAD. Simultaneous co-occurrence has been established to happen between 0.3%-1.2% of cases, and metasynchronous (where secondary cancer develops after the primary one) occurrences happen at about 0.7%-15%^36^, in line with our misclassification rate (1.54%). We therefore suspect that some TCGA lung cancers might be cases of co-occurring adeno and squamous carcinomas.

## Discussion

We have introduced CNSistent, a new Python-based library for processing and exploratory data analysis of SCNA profiles. Using the PCAWG, TCGA, and TRACERx datasets, we have developed a suggested workflow of imputing source data, removing regions of low mappability and filtering samples using z-score based outlier detection. Our conclusion is that it is preferable to filter sample sets from different studies individually before creating a combined dataset, since their individual distributions of CN called regions might significantly differ. In cases where only samples with a certain minimum proportion of SCNAs are of interest, we found GnD as a better threshold criterion than the number of breakpoints.

Using the filtered and combined datasets, we compared several methods of segmentation w.r.t. the task of classification between different cancer types. We observed that a too coarse segmentation (such as chromosome arms) leads to a drop in classification accuracy. To achieve a good balance between accuracy and bias we suggest using 1000-5000 segments based on either chromosome binning or breakpoint clustering. In our case the best solution seemed to be using 2709 one megabase segments. Additionally, we noticed that using the COSMIC gene set performs better than other methods with similar number of segments, suggesting that the CN state of those genes is the driving force of SCNAs selection across cancer types. Finally, we saw that this architecture scaled well to an arbitrary number of sample classes, with 50 classes still achieving a 70% per-class accuracy. Further improvements are to be expected with an increased number of samples.

To investigate the occurrence of batch effects, we compared classification between NSCLC types in all three datasets and saw that the per-sample accuracy improved by combining these three studies when compared to classification of each of them separately. We concluded that when using our workflow, the batch effects are sufficiently small to enable creation of consistent and reliable datasets. We also saw that models trained on one dataset can be successfully applied to classify another, sometimes even outperforming the source dataset, demonstrating that the bias of a classifier toward its training dataset is negligible.

To understand irregularities we observed during the NSCLC classification, we conducted additional interpretability analysis using IGs, combined with exploratory data analysis using the CNSistent plotting tools. We identified *SOX2* gains as under significant positive selection in LUSC where it shows consistent patterns of amplification (average total CN of 7.56). This is combined with negative selection of *EIF4A2*, creating a cliff-shaped profile that is consistently found across samples in all datasets. We concluded that this pattern is both the major classification factor for our network as well as a fundamental feature of this cancer type.

Similar tools to CNSistent have been developed e.g. for RNA-seq datasets^37^, but to the best of our knowledge this is the first such tool for SCNA data. CNSistent closes this gap and provides researchers with a toolkit for the fast and consistent analysis of SCNA profiles from heterogeneous sources. CNSistent enables researchers to start research projects in this field quickly, efficiently analyze copy number data, and develop results in a reproducible and consistent manner.

## Acknowledgements

The results published here are in part based upon data generated by the TCGA Research Network: https://www.cancer.gov/tcga.

The authors would like to thank Tom L. Kaufmann, Cody B. Duncan, Philipp G. Keyl, and Tom B. K. Watkins, Thomas J. Y. Kono, Laura Godfrey, and Daniel Schütte for their feedback.

