## Supplementary Figures 1-9 for "CNSistent integration and feature extraction from somatic copy number profiles"

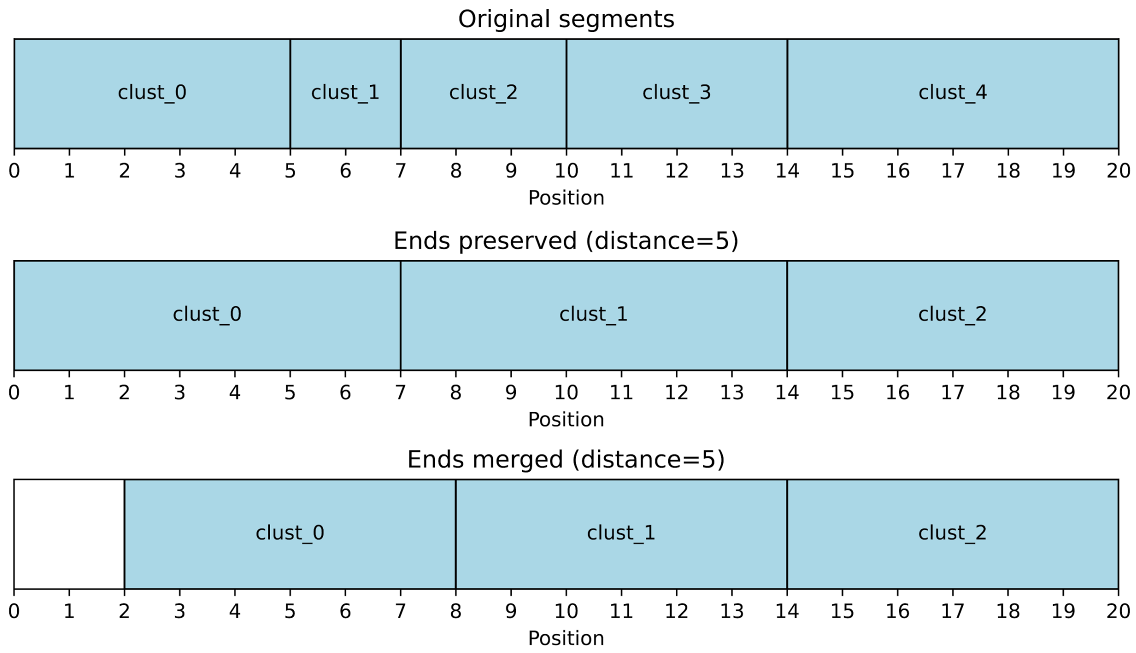


*Supplementary Figure 1: Example of clustering method changing segmentation on a region (top) from 0 to 20, with breakpoints at 5, 7, 10, and 16 . The first version with distance 5 preserves the endpoints, therefore 0 and 20 are not considered; 5, 7, and 10 are merged into 10 (rounding down), while the breakpoint at 14 does not get merged on either side–the previous breakpoint at 10 is closer than 5, but has already been merged with 5 and 7. The second version also merges ends, causing 0 and 5 to merge into 2, while 7 and 10 merge into 8.*

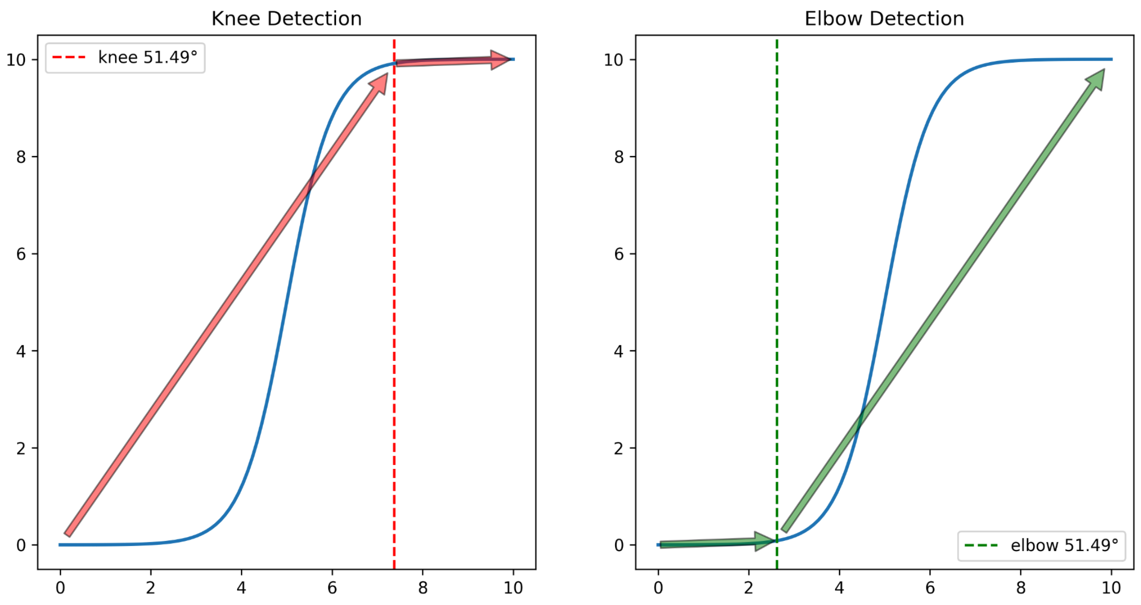


*Supplementary Figure 2: Knee and elbow detection. The angles in the legend are the absolute change in the slope, convex (slope decrease) for the knee, concave (slope increase) for the elbow.*

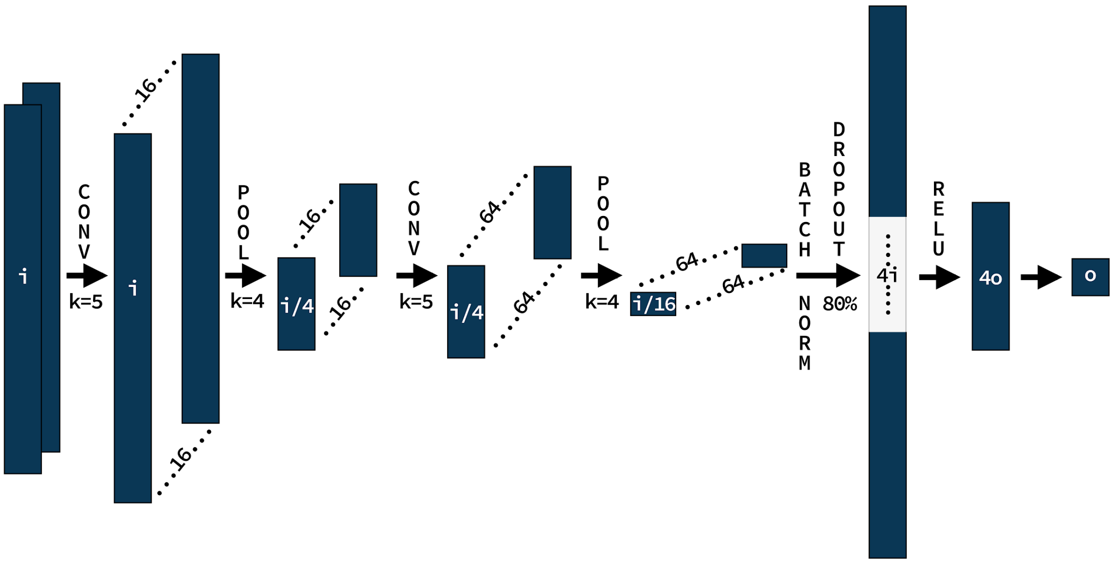


*Supplementary Figure 3: A convolutional neural network classifier with adjustable width depended on the input layer size **i** and the number of classes (output layer size) **o**. We use the Adam optimizer with a learning rate of 0.001 and weight decay of 0.01. To dampen overfitting, we use early stopping when no training loss improvement has been reached in the last 10 epochs.*

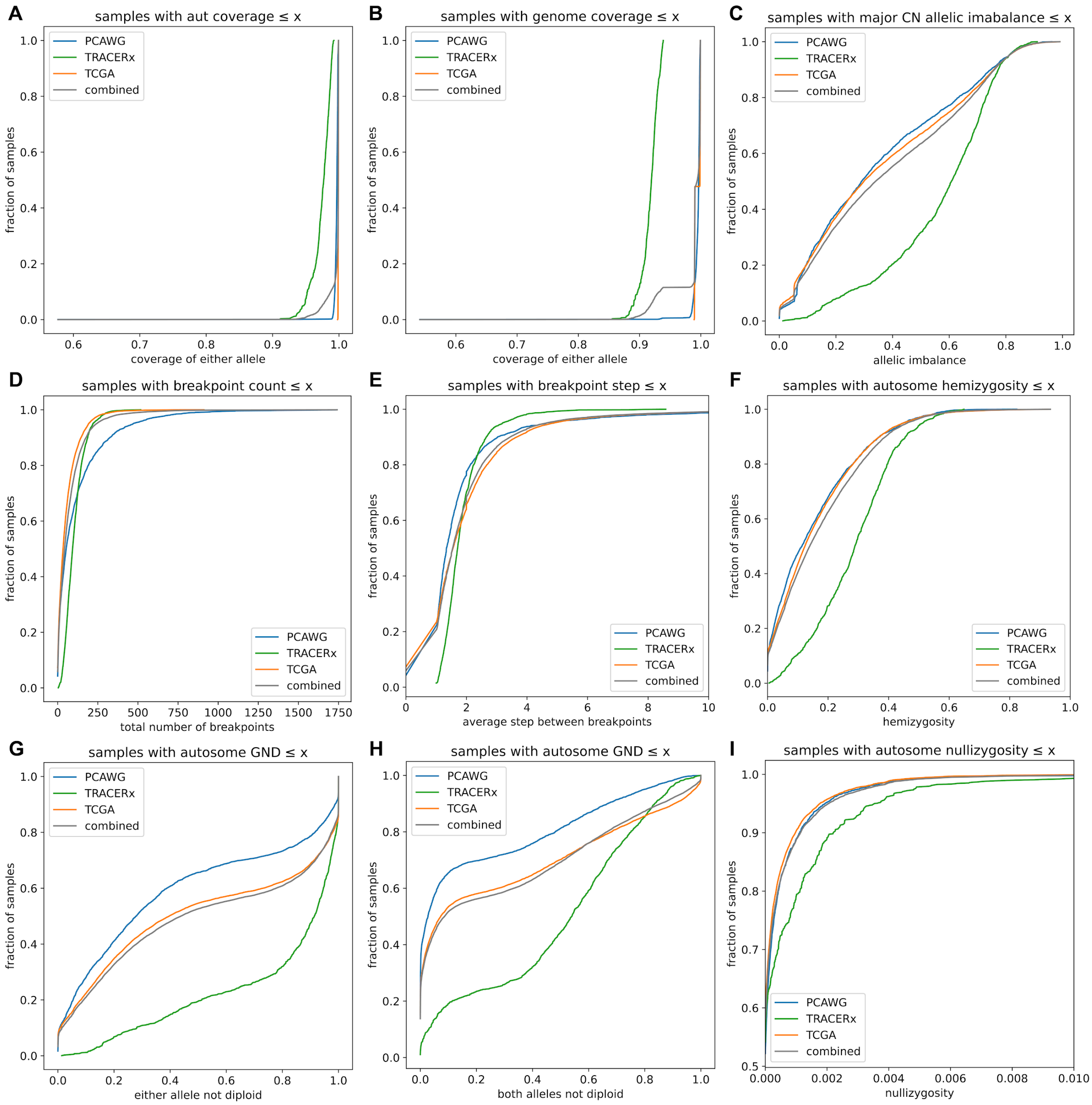


*Supplementary Figure 4: Summary features across datasets. Unless otherwise specified, only autosomes are considered. Except for D and E, all features are calculated using gap-mask.* ***A)*** *Coverage for autosomes only,* ***B)*** *Coverage of the whole genome - note the shift on the TRACERx samples which lack sex chromosomes.* ***C)*** *Proportion of the genome with the major allele has higher CN than the minor one.* ***D)*** *Number of breakpoints per sample.* ***E)*** *The average step per breakpoint - there is no value between 0 and 1 as any two breakpoints differ by at least 1.* ***F)*** *Proportion of samples with one of the alleles lost.* ***G)*** *Genome not diploid on either of the alleles in the autosomes.* ***H)*** *Genome not diploid on both of the alleles in the autosomes.* ***I)*** *Proportion of samples where both alleles are 0. The x-axis is scaled for better visibility, as barely more than 1% of samples exhibit nullizygosity.*

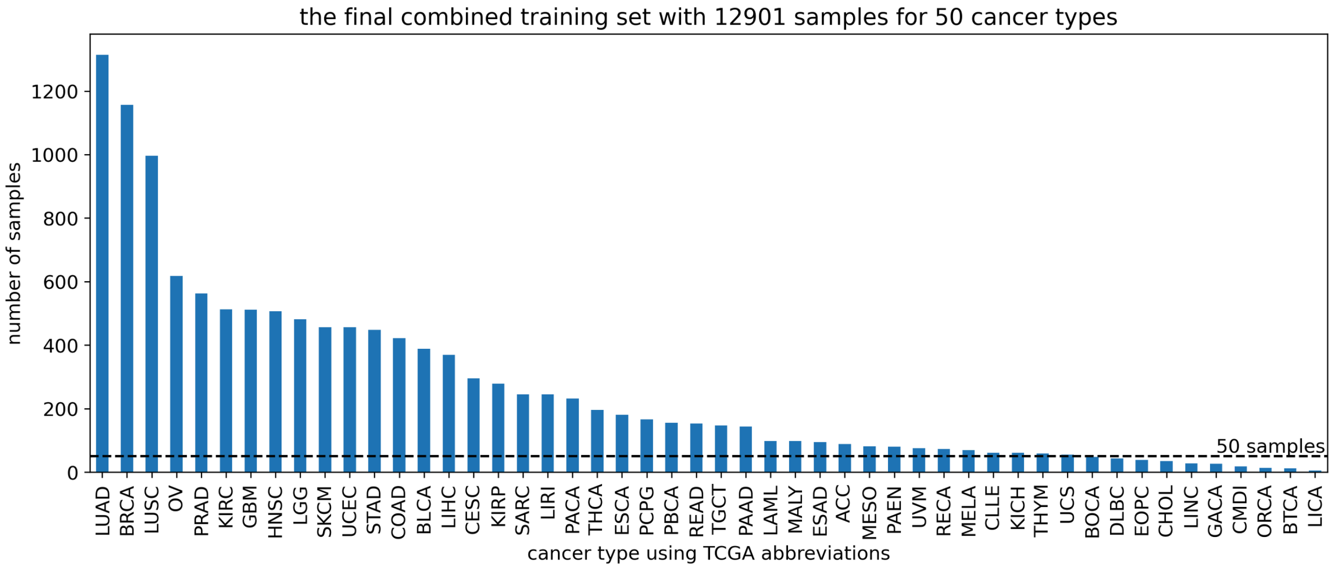


*Supplementary Figure 5: Distribution of samples per cancer type in the final joint dataset.*

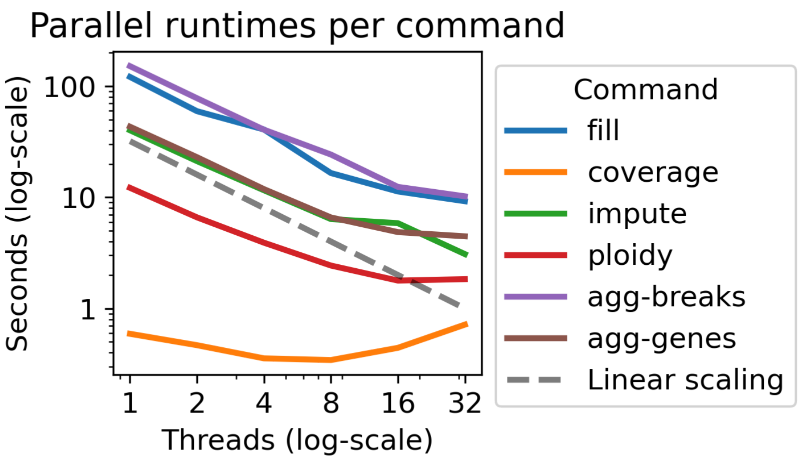


*Supplementary Figure 6: Runtime of individual CNSistent commands across 1, 2, 4, 8, 16, and 32 threads show near-linear scaling on the PCAWG dataset.*

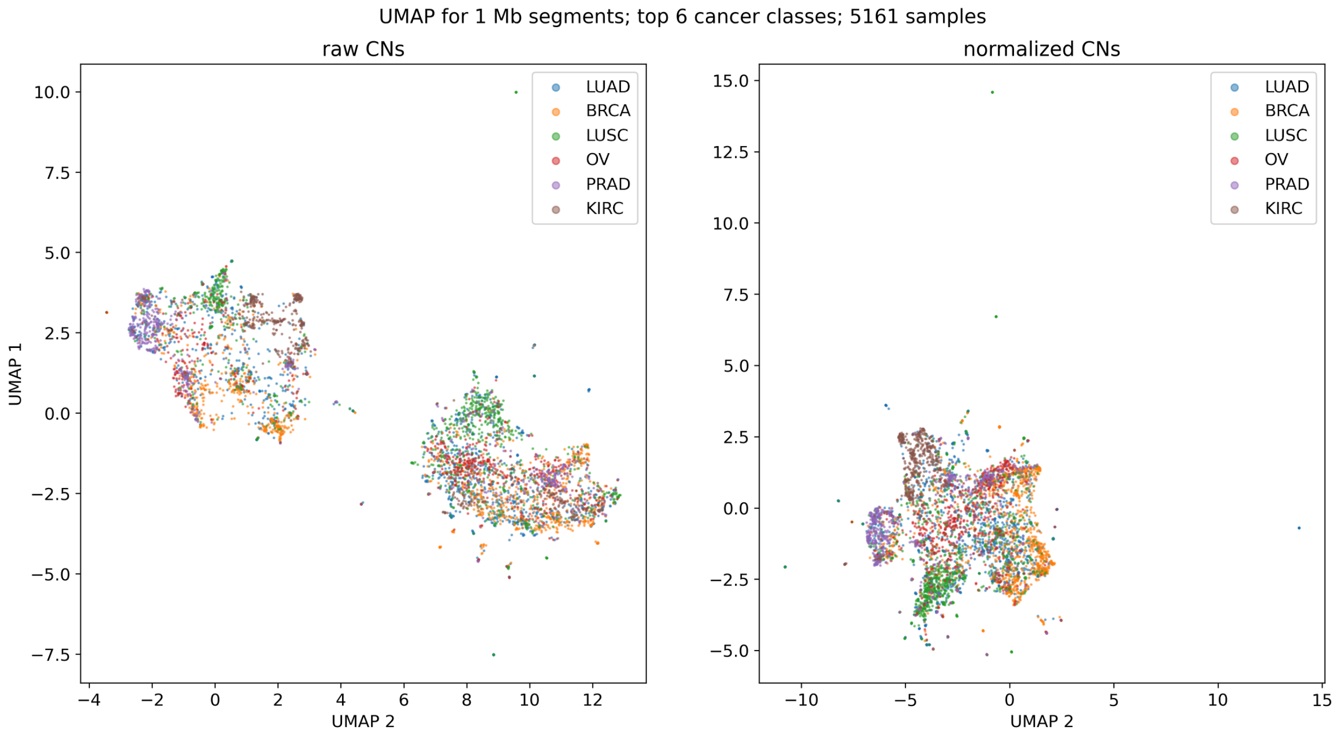


*Supplementary Figure 7: A UMAP plot for the top 6 classes in the combined dataset using 1 Mb segmentation with gaps removed. The minor and major CNs are concatenated into a single input vector per sample. We can see a certain level of clustering per type, however clear separation would be difficult. In the left plot, original values are used, creating two distinct clusters, presumably separating the samples with and without whole genome doubling. In the right plot one we normalize the CNs so that the mean CN of each sample is 1, creating a single cluster. UMAP.fit() function from the umap-learn package version 0.5.6 has been used with default parameters. Note that despite there being always multiple samples from the same patient, there is no clear aggregation.*

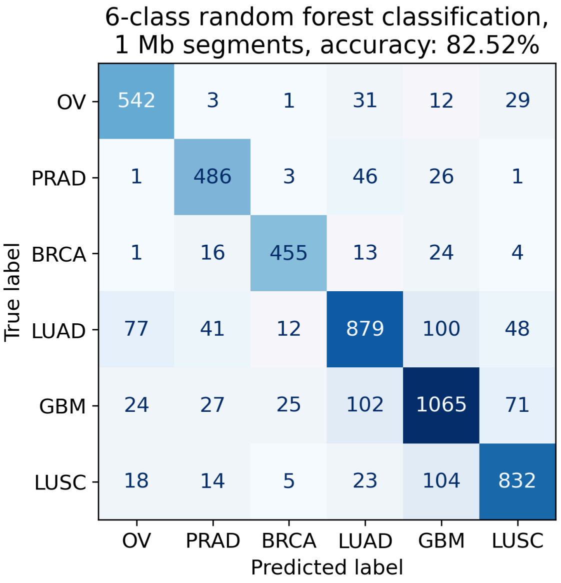


*Supplementary Figure 8: Random forest classifier of the top 6 cancer types yields 82.52% accuracy using the 1 Mb segmentation with gaps removed. A 5-fold cross-validation using the RandomForestClassifier object from scikit-learn 1.2.2. with default parameters has been used.*

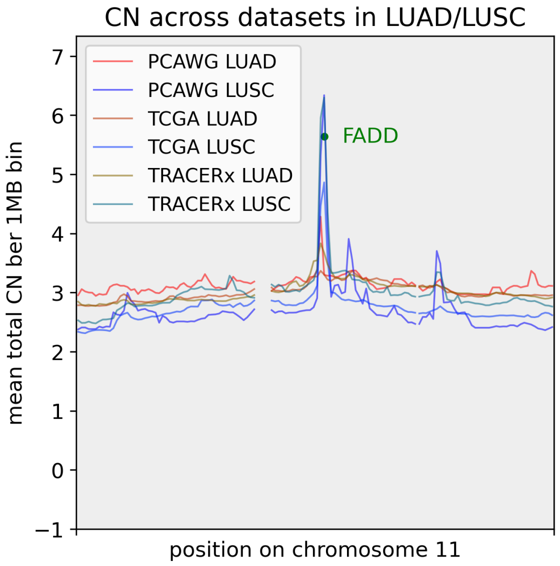


*Supplementary Figure 9: CN profiles of LUNG/LUSC on chromosome 11. There is a considerable spike at the location of the gene FADD which is strongly overexpressed in LUSC, with the average CN of 5.64.*
